# Rieske FeS overexpression in tobacco provides increased abundance and activity of Cytochrome *b*_6_*f*

**DOI:** 10.1101/2022.06.28.497970

**Authors:** Eiri Heyno, Maria Ermakova, Patricia E. Lopez-Calcagno, Russell Woodford, Kenny L. Brown, Jack S. A. Matthews, Barry Osmond, Christine A. Raines, Susanne von Caemmerer

## Abstract

Photosynthesis is fundamental for plant growth and yield. The Cytochrome *b*_6_*f* complex catalyses a rate-limiting step in thylakoid electron transport and therefore represents an important point of regulation of photosynthesis. Here we show that overexpression of a single core subunit of Cytochrome *b*_6_*f*, the Rieske FeS protein, led to up to a 40% increase in the abundance of the complex in *Nicotiana tabacum* (tobacco) and was accompanied by an enhanced *in vitro* Cytochrome *f* activity, indicating a full functionality of the complex. Analysis of transgenic plants overexpressing Rieske FeS by the light-induced fluorescence transients technique revealed a more oxidised primary quinone acceptor of Photosystem II (Q_A_) and plastoquinone pool and a faster electron transport from the plastoquinone pool to Photosystem I upon changes in irradiance, compared to control plants. A faster establishing of q_E_, the energy-dependent component of non-photochemical quenching, in transgenic plants suggested a more rapid build-up of the transmembrane proton gradient, also supporting the increased *in vivo* Cytochrome *b*_6_*f* activity. However, there was no consistent increase in steady-state rates of electron transport or CO_2_ assimilation in plants overexpressing Rieske FeS grown in either laboratory conditions or in field trials, suggesting that the *in vivo* activity of the complex was only transiently increased upon changes in irradiance. Our results show that overexpression of Rieske FeS in tobacco enhances abundance of functional Cytochrome *b*_6_*f* and electron transport capacity and may have a potential to increase plant productivity if combined with other traits.

**One-sentence summary:** Increased abundance of Cytochrome *b*_6_*f* complex leads to transient increases in photosynthetic electron transport rate in tobacco.

## Introduction

Global crop production needs to double by 2050 to meet the needs of increasing human and cattle populations and biofuel consumption (Ray et al., 2013). The green revolution has brought many parameters of crop yield potential close to the theoretical maximum, the most significant ones being the efficiency of biomass partitioning into the harvested product and efficiency of solar radiation capture. Modelling studies suggest that there is still significant potential in increasing the efficiency of photosynthesis, the major contributor to plant growth (Long et al., 2006; Evans, 2013; Parry et al., 2013; Ort et al., 2015; Bailey-Serres et al., 2019). This has led to research efforts focusing on improving various photosynthetic components in attempt to improve plant productivity (Kromdijk et al., 2016; López-Calcagno et al., 2019; Ermakova et al., 2021).

Photosynthetic electron transport is responsible for the provision of ATP and NADPH for photosynthetic carbon reduction and it is suggested to limit CO_2_ assimilation rates at ambient and high CO_2_ partial pressures (von Caemmerer and Farquhar, 1981; Yamori et al., 2010). Light energy, captured by light-harvesting pigments, drives electron transport through the thylakoid membranes of chloroplasts. The Cytochrome *b*_*6*_*f* complex (Cyt*b*_*6*_*f*) plays a central role in the electron transport chain as it functions as an electron hub between Photosystem II (PSII) and Photosystem I (PSI), and its proton pumping activity is the main contributor to the acidification of the thylakoid lumen. Cyt*b*_*6*_*f* also has a unique role in electron transport because it is involved in both linear electron transport, carried out by both photosystems to produce ATP and NADPH, and in cyclic electron transport around PSI for the generation of ATP alone. Transmembrane proton gradient (ΔpH) established across the thylakoid membrane during the electron transport is the main component of the proton motive force (*pmf*) used by the ATP synthase complex to produce ATP. In addition, ΔpH exerts control over the rate of electron transport by a feedback regulation of PSII activity via non-photochemical quenching (NPQ) that dissipates a part of light energy absorbed in the PSII antennae as heat before it reaches the reaction centres (Ruban, 2016). The major, quickly reversible component of NPQ (q_E_) is mediated by the PsbS protein acting as a luminal pH sensor and by violaxanthin de-epoxidase converting violaxanthin to zeaxanthin in the PSII antennae (Johnson et al., 2009).

Cyt*b*_*6*_*f* catalyses the slowest reaction in the electron transport chain (Tikhonov, 2014). Moreover, acidification of the lumen further slows down the Cyt*b*_*6*_*f* activity, a phenomenon also known as photosynthetic control (Colombo et al., 2016). Previous works on transgenic tobacco plants with reduced abundance of the Rieske FeS protein (hereafter Rieske), one of the core subunits of Cyt*b*_*6*_*f* encoded by the nuclear *petC* gene, demonstrated a strong correlation between the amount of the complex in the thylakoid membrane and leaf electron transport rate (Price et al., 1995; Price et al., 1998; Ruuska et al., 2000; Yamori et al., 2011). Similar results were observed in rice, where plants with reduced Cyt*b*_*6*_*f* content showed decreased grain yield (Yamori et al., 2016). Overexpression of Rieske in the model C_3_ and C_4_ species, *Arabidopsis thaliana* and *Setaria viridis*, led to increases in electron transport and CO_2_ assimilation rates, suggesting that it is a promising route to boost crop productivity (Simkin et al., 2017; Ermakova et al., 2019).

Here we explored the effect of Rieske overexpression in the model crop plant *Nicotiana tabacum* (tobacco). We show that increasing Rieske content provides up to 20% increase in functional Cyt*b*_6_*f*, as confirmed by the Cytochrome *f* (CytF) activity, but without marked changes in steady-state electron transport or CO_2_ assimilation rates. Our detailed functional analysis of electron transport using light-induced fluorescence transients technique and spectroscopic measurements confirmed higher Cyt*b*_*6*_*f* activity in plants with increased Rieske content. Our results indicate that, at tested CO_2_ and irradiance regimes, Cyt*b*_*6*_*f* activity is not the only factor limiting electron transport rate in tobacco.

## Results

### Generation of transgenic tobacco plants

Using *Agrobacterium*-mediated leaf transformation, *N. tabacum* was transformed with a construct for Rieske overexpression (Rieske-OE hereafter, Fig. S1). From 20 independent lines generated, 9 lines with a single gene insertion were screened for transgene expression, and transcripts of the transgenic *petC* were detected in 8 of those lines in T_1_ generation (Fig. 1). Homozygous plants were selected from the T_1_ progenies of lines R13, R17, R20, R25 and R26. The progenies of the homozygous plants were used in all further analyses.

**Fig. 1.**
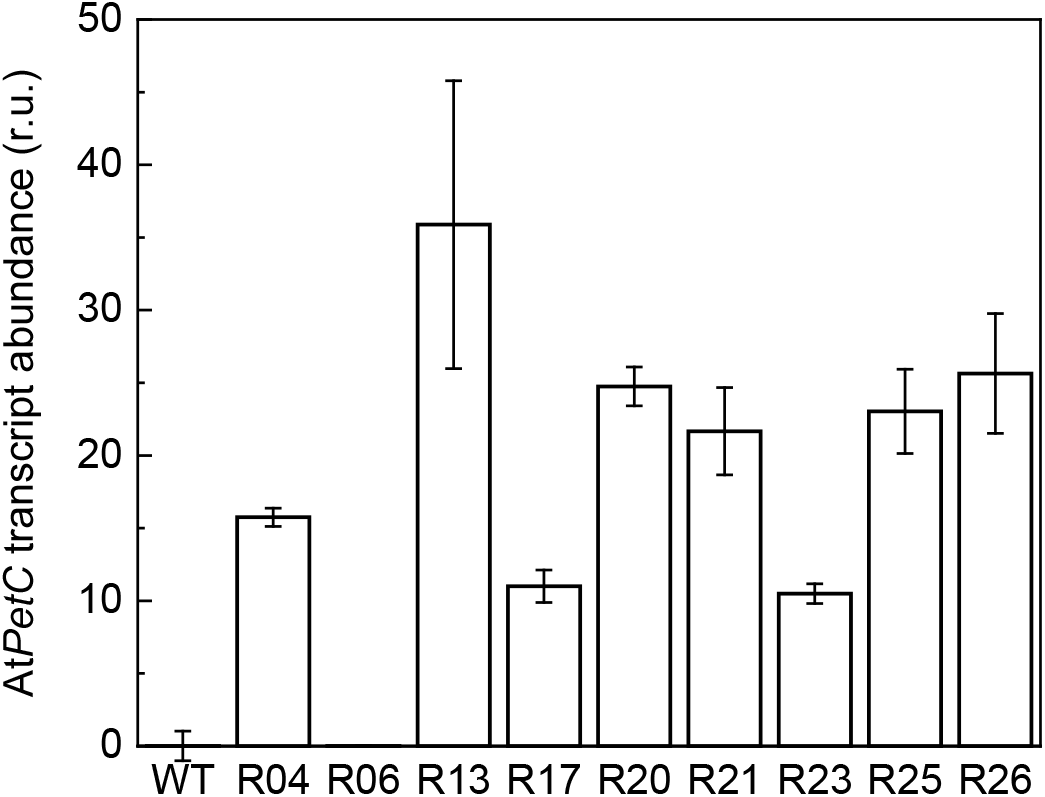
Transcript abundance of *AtPetC* in T_1_ plants compared to WT. Relative *AtPetC* expression as fold of *NtActin* in homozygous T_1_ plants and WT plants. The individuals used in this analysis generated the T_2_ seeds used for the phenotyping experiments presented below. Mean ± SD, *n* = 3 technical replicates.

### Cytochrome b_6_f is functional in Rieske-OE plants

Rieske-OE plants were analysed for Cyt*b*_6_*f* abundance and activity. To estimate the content of Cyt*b*_6_*f*, we separated thylakoid protein complexes from wild type (WT) and Rieske-OE plants by Blue Native-PAGE and detected the complex using Rieske antibody (Fig. 2, Fig. S2). A band matching the size of a dimeric Cyt*b*_6_*f* complex of approximately 400 kDa was recognised (Kügler et al., 1997). Relative quantification showed a 20-60% increased abundance of Cyt*b*_6_*f* in all lines (significant in lines R13 and R17) (Fig. 2, Fig. S2). To verify the functionality of the complex in Rieske-OE plants, we used the reduced-oxidised (hydroquinone minus ferricyanide) difference spectroscopy assay of CytF, one of the redox cofactors of Cyt*b*_6_*f*. The absorption peak of the characteristic alpha band at 554 nm was used to quantify the amount of CytF in the sample using the extinction coefficient of 17.7 cm^2^ mmol^-1^ (Fig. 3a). All lines, except for R13, displayed a 10-30% increase of CytF activity, significant for lines R17, R20 and R26 (Fig. 3b). WT CytF activity remained constant between different batches of plants (Fig. S3).

**Fig. 2.**
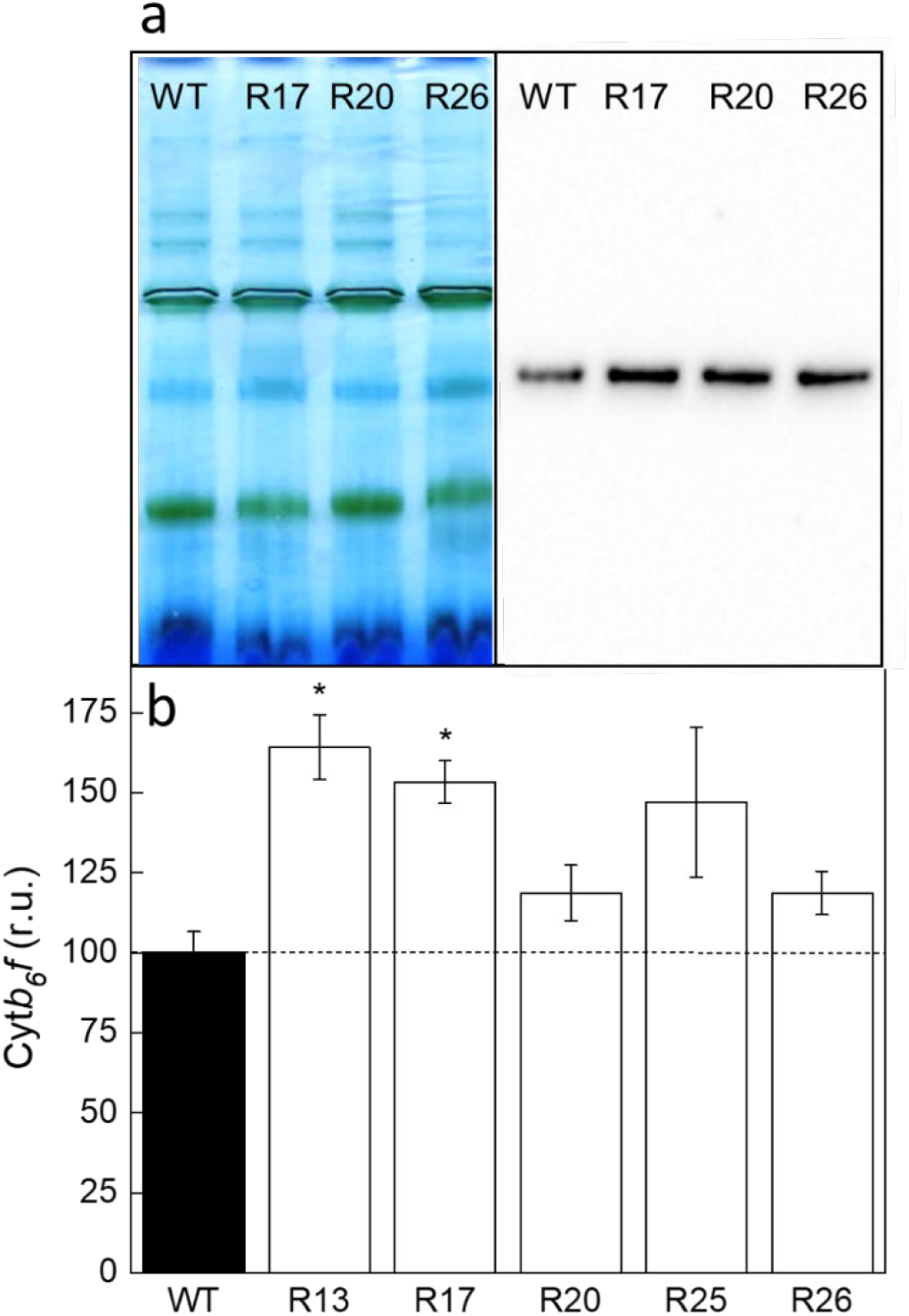
Immunodetection of Cytochrome *b*_6_*f* complex from the thylakoid membranes of WT and Rieske-OE plants. (**a**) Blue Native-PAGE of the thylakoid protein complexed followed by the western blotting with Rieske antibodies (left and right panels, respectively). 10 µg Chl (*a*+*b*) of each sample was loaded. Additional gel and blot images of all lines tested are shown in Fig. S2. (**b**) Relative quantification of the blots. Mean ± SE, *n* = 3 biological replicates. Asterisks indicate statistically significant differences between transgenic lines and WT (*t*-test, *P* < 0.05).

**Fig. 3.**
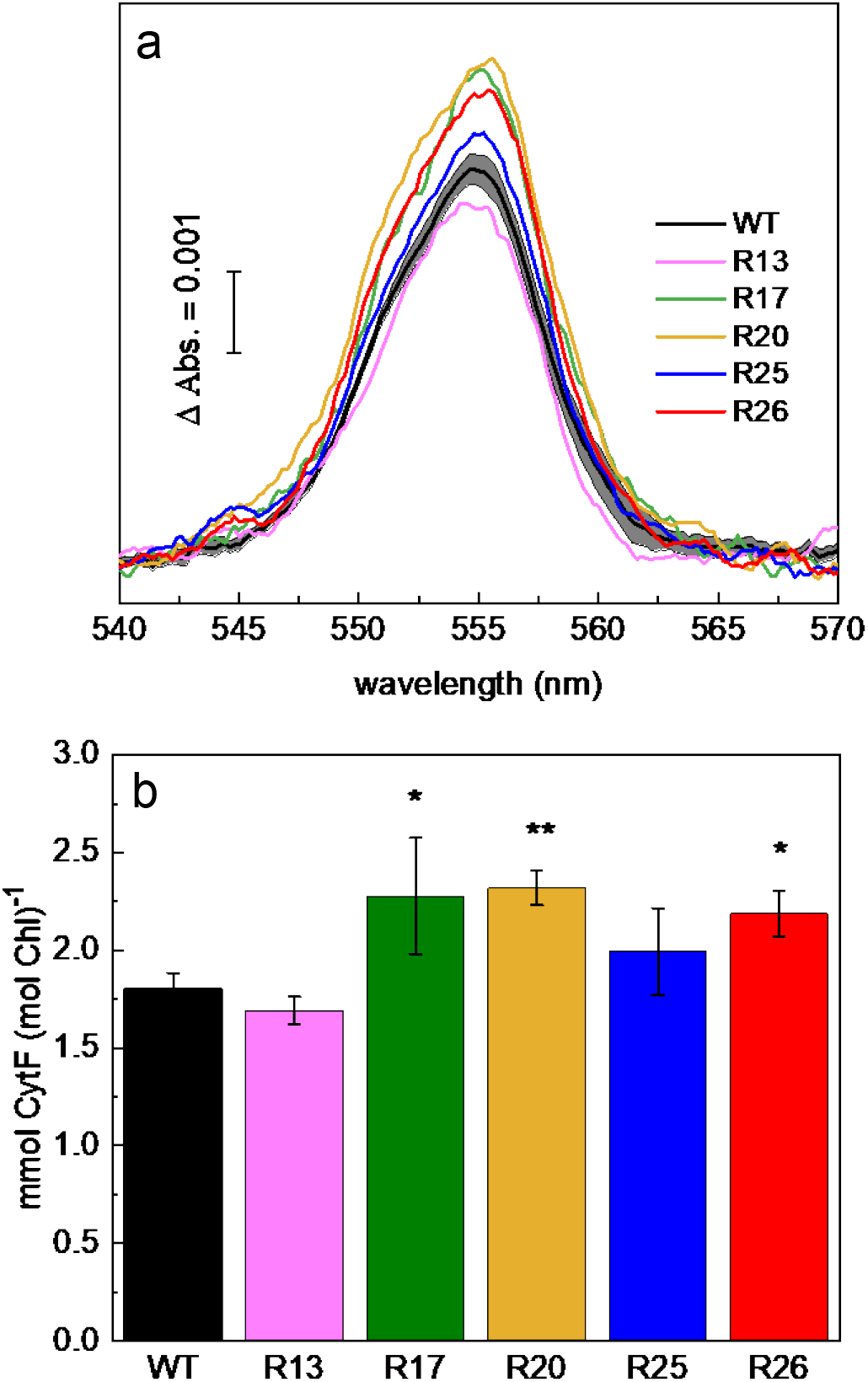
Quantification of CytF content in the thylakoid membranes of WT and Rieske-OE plants. (**a**) The reduced-minus-oxidised CytF spectra. Traces are average of 3-8 biological replicates; SE range is shown for the WT trace. (**b**) Relative abundance of CytF calculated from the 554 nm peak. Mean ± SE, *n* = 8 biological replicates for WT, *n* = 3 for transgenic lines. Asterisks indicate statistically significant differences between transgenic lines and WT (*t*-test, \**P* < 0.05, \*\**P* < 0.01).

Next, we selected lines R13, R17, R25 and R26 to analyse the abundance of photosynthetic proteins in leaf extracts by immunoblotting with specific antibodies (Fig. 4). A significant, near 2-fold increase of Rieske abundance was confirmed in all lines on leaf area basis. No changes in abundance of D1 (PSII), PsaB (PSI) nor AtpB (ATP synthase) were detected in any transgenic lines, suggesting an unaltered abundance of the corresponding thylakoid complexes. The content of the large subunit of Rubisco (RbcL) and PsbS did not differ between the WT and Rieske-OE plants. Interestingly, the soluble electron carrier between Cyt*b*_6_*f* and PSI, plastocyanin (PC), and the Lhcb2 subunit of the light-harvesting complex II (LHCII) displayed an opposite trend between the lines, with a significant decrease of PC in lines R25 and R26 and a significant decrease of Lhcb2 in lines R13 and R17.

**Fig. 4.**
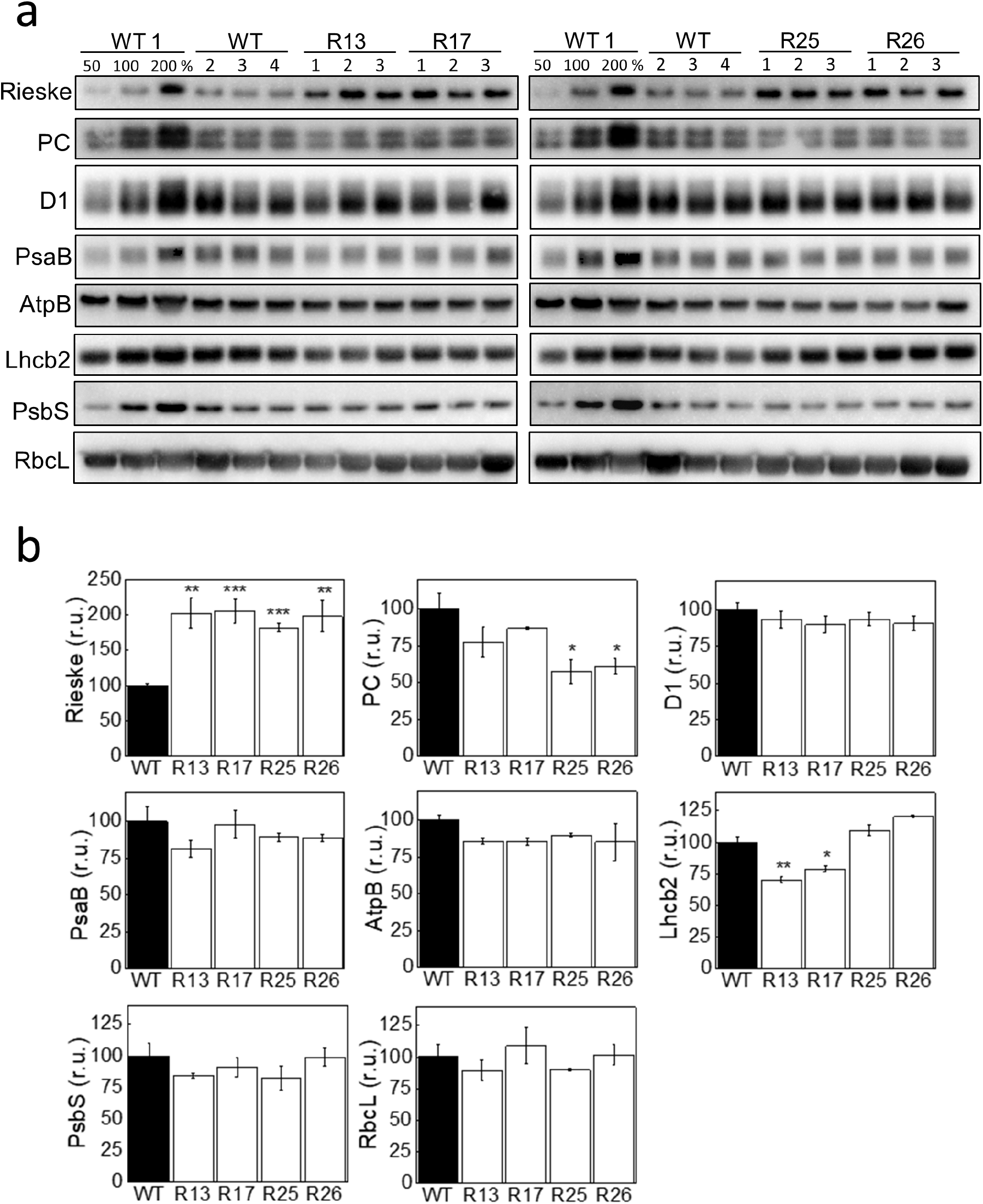
Abundance of photosynthetic proteins in leaves of WT and Rieske-OE plants. (**a**) Immunodetection of Rieske, Plastocyanin (PC), D1 subunit of PSII, PsaB subunit of PSI, AtpB subunit of ATP synthase, Lhcb2 subunit of LHCII, PsbS and the large subunit of Rubisco (RbcL) in leaf protein extracts loaded on leaf area basis. A titration series of one of the WT samples (WT1) was used for relative quantification. (**b**) Quantification of immunoblots relative to WT. Mean ± SE, *n* = 4 biological replicates for WT, *n* = 3 for transgenic lines. Asterisks indicate statistically significant differences between transgenic lines and WT (*t*-test, \**P* < 0.05, \*\**P* < 0.01).

### Overexpression of Rieske changes light-induced fluorescence transients

To analyse the function of Cyt*b*_6_*f* in Rieske-OE plants *in vivo*, we studied electron transport rates by the light-induced fluorescence transients (LIFT) technique using the fast repetition rate (FRR) method (Kolber et al., 1998). This near-remote sensing method (0.6-40 m) is finding increasing applications in phenotyping of photosynthetic electron transport kinetics in the field (Keller et al., 2019; Keller et al., 2019). The advantages of this method are fast time resolution and minimally intrusive *in situ* approach via the use of sub-saturating short excitation pulses (‘flashlets’) that exert minimal changes to the redox state of electron carriers as opposed to high intensity flashes used in pulse amplitude modulation (PAM) fluorescence technique. Using recently developed LIFT protocols for laboratory analysis of plants (Osmond et al., 2019; Osmond et al., 2021), we assessed *in vivo* activity of Cyt*b*_6_*f* in tobacco. LIFT-FRR method allows for real-time monitoring of F_V_/F_M_ (reporting on the redox states of Q_A_, the primary quinone acceptor of PSII), the redox state of the plastoquinone (PQ) pool and electron transport rates from PSII to PQ pool (Tau1) and from PQ pool to PSI (Tau2) (Osmond et al., 2019; see also under Photosynthetic signatures at https://soliense.com/LIFT_Method.php). The Q_A_ is first gradually reduced by flashlets at microsecond intervals until the fluorescence signal is saturated (300 flashlets). This is immediately followed by a Q_A_ re-oxidation flash routine, during which the time between the flashlets is exponentially increased to allow for graduate re-oxidation (90 flashlets). Each flashlet cycle, *i*.*e*., the excitation and re-oxidation flashlets (390 in total), yields to ∼300 ms chlorophyll (Chl) fluorescence transients (Fig. S4) that are fitted by the FRR model to derive Chl fluorescence parameters.

Illumination of the dark-adapted tobacco leaves with far-red light caused an increase in Tau1, indicating a slower electron transport from PSII to the PQ pool, as well as a more oxidised PQ pool and a decrease in Tau2, indicating faster electron transport from the PQ pool to PSI (Fig. 5). The effect of far-red on Tau1 remained throughout the FR illumination whilst Tau2 gradually returned to the level observed in darkness, showing a slow-down of electron transport to PSI. These observations were consistent with an operation of far-red light-driven cyclic electron flow and showed no difference between the WT and transgenic plants. Next, red actinic light of 400 µmol m^-2^ s^-1^ was added on top of the far-red illumination (indicated by arrows in Fig. 5), which induced a rapid reduction of Q_A_ (seen as a drop in F_V_/F_M_) and PQ pool followed by a gradual re-oxidation until it reached a quasi-steady-state level after approximately three minutes. Tau1 and Tau2 displayed reproducible transients with one or two peaks (*i*.*e*., slower electron transport), respectively, before reaching a steady-state. Line R17 displayed significantly decreased Tau1 and Tau2 values, corresponding to faster electron transport rates. Line R26 also had decreased Tau2. Line R25 showed a similar trend of F_V_/F_M_ as the other lines, although not significant, but the Tau2 values were significantly lower in this line. Interestingly, initial differences in Q_A_ and PQ pool oxidation between WT and transgenic lines, seen in the beginning of actinic light illumination, were often reduced by the end of the illumination period. Collectively, these results suggested that Rieske-OE lines maintained transiently more oxidised PQ pool and Q_A_ upon changes in irradiance, likely, due to the increased Cyt*b*_6_*f* activity.

**Fig. 5.**
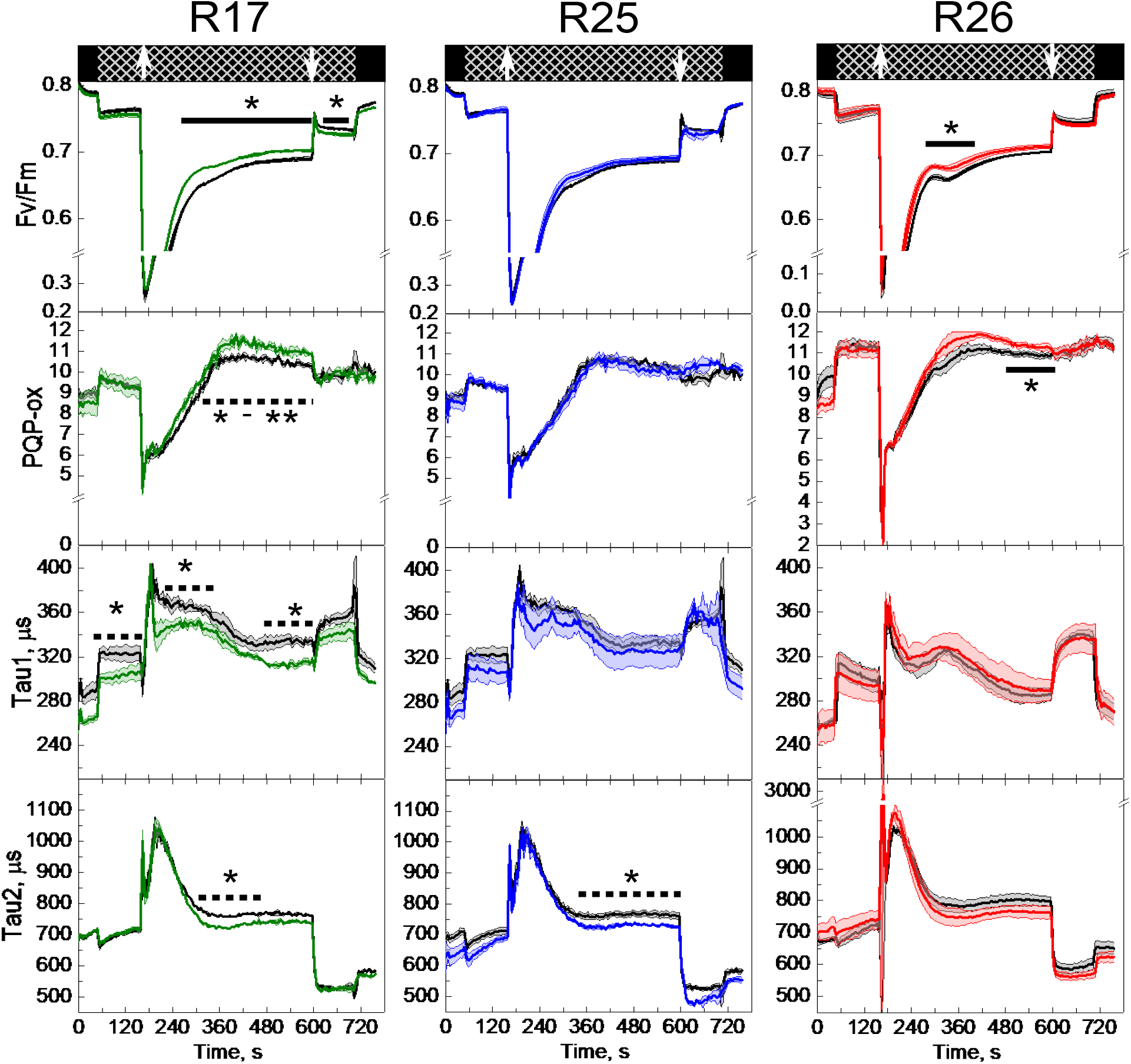
Electron transport parameters of WT (black traces) and Rieske-OE lines R17, R25 and R26 derived from the light-induced fluorescent transients measurements. The upper panels indicate background illumination during the measurement: black = darkness, hatched = far-red light of 200 µmol m^-2^ s^-1^, arrows indicate the onset and offset of red actinic light of 400 µmol m^-2^ s^-1^. F_V_/F_M_, oxidised Q_A_ sites; PQP_ox, oxidation state of the plastoquinone pool; Tau1, electron transport rate from PSII to the plastoquinone pool; Tau2, electron transport rate from the plastoquinone pool to PSI. Mean ± SE, *n* = 3 biological replicates for transgenic lines, *n* = 4 for WT. Asterisks indicate statistically significant differences between transgenic lines and WT over a time period (*t*-test, \**P* < 0.05, \*\**P* < 0.01). Broken line indicates that at least 50% of the data points were significant.

### Thylakoid membrane energisation in Rieske-OE plants

Next, we assessed *pmf* established in leaves of WT and Rieske-OE plants at different irradiance by monitoring the dark interval relaxation kinetics of the electrochromic shift signal (Fig. 6a). Line R26 showed significantly increased *pmf* at 1000 µmol m^-2^ s^-1^ and above whilst lines R17 and R25 did not differ from the WT. No differences in the proton conductivity of the thylakoid membrane (*g*_H+_) were observed between the genotypes suggesting that activity of the chloroplast ATP synthase was not altered in Rieske-OE plants (Fig. 6b). Absorption changes at 535 nm upon the illumination largely account for the q_E_ formation in response to acidification of the lumen (Wilson et al., 2021). Upon the shift from dark to 400 µmol m^-2^ s^-1^, all three measured transgenic lines established q_E_ more rapidly than WT (Fig. 6c) consistent with a faster build-up of ΔpH in plants with increased Cyt*b*_6_*f* activity.

**Fig. 6.**
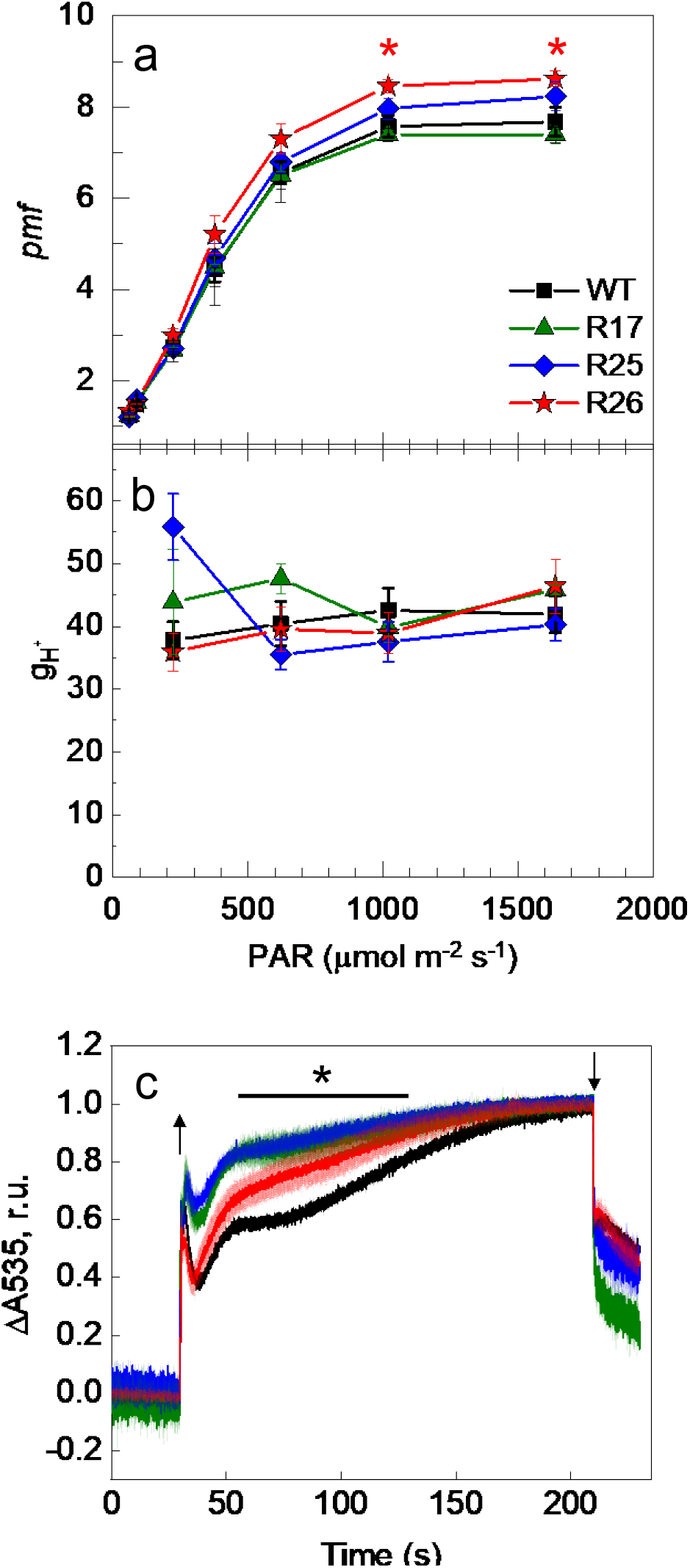
Analysis of the thylakoid membrane energisation in WT and Rieske-OE plants. (**a** and **b**) Proton motive force (*pmf*) and proton conductivity of the thylakoid membrane (*g*_H_^+^) at different irradiances. Mean ± SE, *n* = 4 biological replicates. (**c**) Changes in absorbance at 535 nm during the dark-light-dark transitions. Traces were normalised to 0 in the beginning of illumination (400 μmol m^-2^ s^-1^, arrow up) and to 1 in the end of illumination (arrow down) to facilitate comparison of the kinetics. Traces are average of 3 biological replicates, SE is shown as a lighter shade. Asterisks indicate statistically significant differences between transgenic lines and WT (*t*-test, *P* < 0.05).

### Gas exchange and electron transport rates of Rieske-OE plants

We studied effects of increased Cyt*b*_6_*f* activity in tobacco on the net CO_2_ assimilation rate and effective quantum yield of PSII, Y(II), at different CO_2_ partial pressures. No significant changes in either parameter were observed in Rieske-OE lines, compared to WT (Fig. 7). The maximum carboxylation rate allowed by Rubisco (*V*_cmax_) and the rate of photosynthetic electron transport based on NADPH requirement (*J*) obtained by fitting the CO_2_ response curves of CO_2_ assimilation showed no changes between the genotypes (Table 1). The leaf mass per area and total leaf Chl content were not altered in the transgenic lines, compared to WT (Table 1). There was a slight increase in the Chl *a*/*b* ratio in lines R13 and R17, which could reflect the changes in the Lhcb2 amounts seen in those lines (Fig. 4).

**Table 1.**
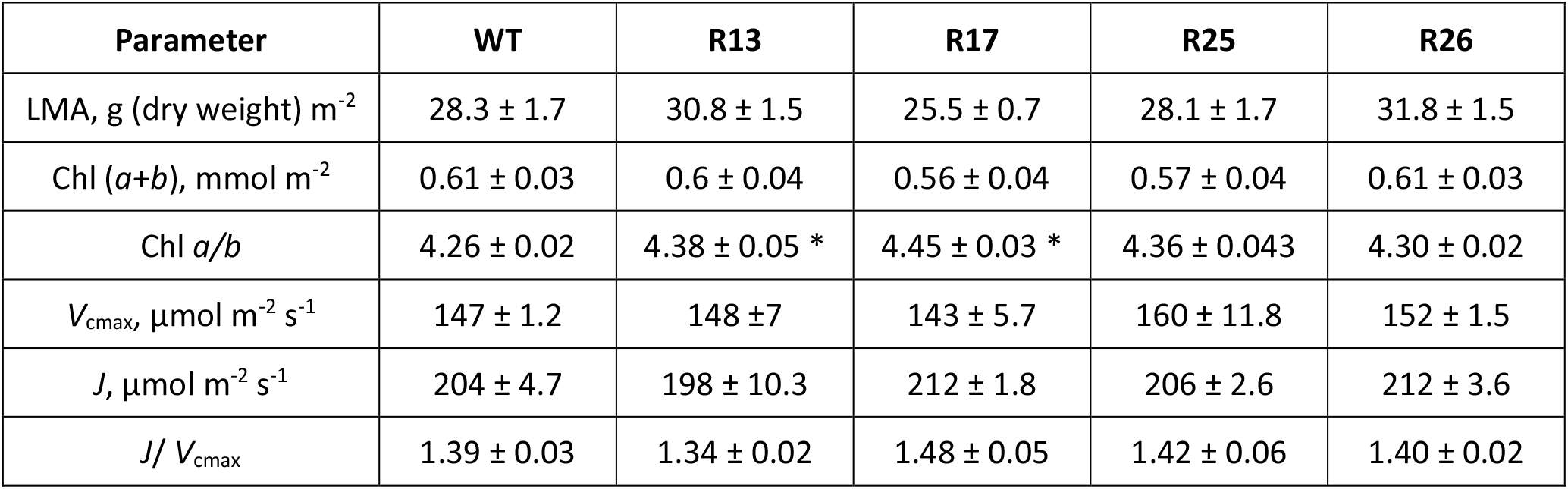
Gas exchange and biochemical properties of WT and Rieske-OE tobacco plants. LMA, leaf mass per area; *V*_cmax_, maximum carboxylation rate allowed by Rubisco; *J*, rate of electron transport based on NADPH requirement. Samples were collected from the leaf area used for gas exchange measurements presented in Fig. 7. Mean ± SE, *n* = 3 biological replicates. Asterisks indicate statistically significant differences between transgenic lines and WT (*t*-test, *P* < 0.05).

**Fig. 7.**
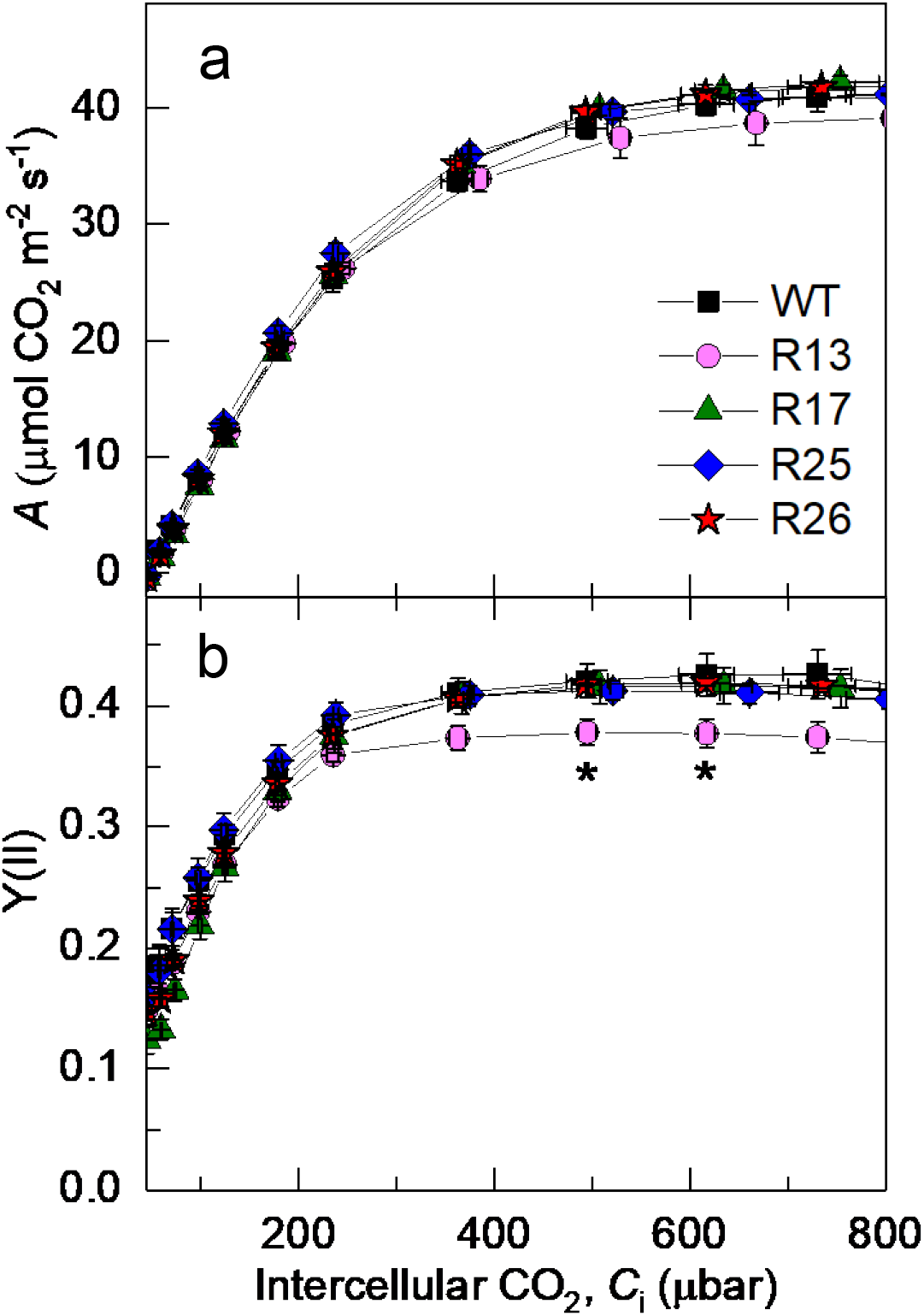
Photosynthetic properties of WT and Rieske-OE tobacco plants grown in controlled conditions. (**a**) CO_2_ assimilation rate, *A*; (**b**) the effective quantum yield of PSII, Y(II). Measurements were performed at 1500 µmol m^-2^ s^-1^ and different CO_2_ partial pressures. Mean ± SE, *n* = 3 biological replicates for Rieske-OE lines, *n* = 6 for WT. Asterisks indicate statistically significant differences between transgenic lines and WT (*t*-test, *P* < 0.05).

### Field trials

To study effects of increased Cyt*b*_6_*f* activity on CO_2_ assimilation and electron transport in tobacco grown in field conditions, three Rieske-OE lines and control plants were grown to maturity in two field sites in Puerto Rico and Illinois (the randomized design of field trails is shown in Fig. S5). When grown in Puerto Rico, all Rieske-OE lines showed significantly increased CO_2_ assimilation rates, compared to control plants, at light intensities above 900 µmol m^-2^ s^-1^ (Fig. 8, left panels). Transgenic plants had similar Y(II) to control plants but showed higher *C*_i_/*C*_a_ (the ratio of intercellular to ambient CO_2_ partial pressures) above 900 µmol m^-2^ s^-1^, suggesting that the detected increases in assimilation were not related to improvements in electron transport, but differences in stomatal conductance. The Illinois field experiments revealed no changes between the genotypes in CO_2_ assimilation, Y(II) or *C*_i_/*C*_a_ at any irradiance (Fig. 8, right panels). CO_2_ responses of CO_2_ assimilation and Y(II) did not differ between transgenic and control plants grown in Illinois (Fig. S6).

**Fig. 8.**
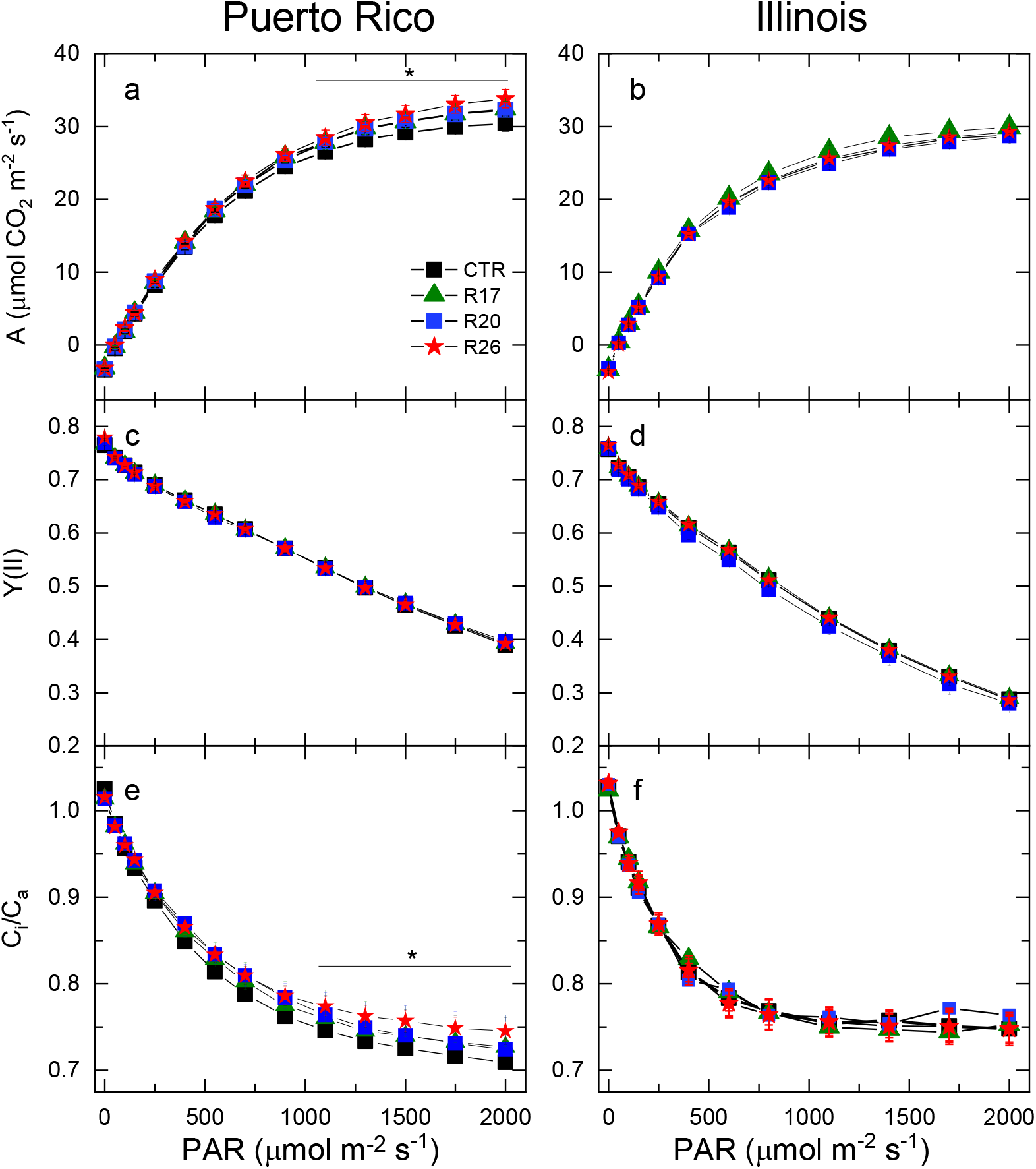
Light responses of photosynthetic parameters measured at ambient CO_2_ from control and Rieske-OE tobacco plants grown in the field. (**a, c, e**) Puerto Rico trials; (**b, d, f**) Illinois field trials. (**a** and **b**) CO_2_ assimilation rate, *A*; (**c** and **d**) the effective quantum yield of PSII, Y(II); (**e** and **f**) the ratio of intercellular and ambient CO_2_ partial pressures, *C*_i_/*C*_a_. The control group (CTR) contained WT and azygous plants. Mean ± SE, *n* = 18 biological replicates for CTR in Puerto Rico, *n* = 24 for CTR in Illinois, *n* = 9 for transgenic lines in Puerto Rico, *n* = 12 for transgenic lines in Illinois. Asterisks indicate statistically significant differences between transgenic lines and WT (type II ANOVA, *P* < 0.05).

## Discussion

Improving photosynthesis is one of the promising routes to increase biomass and yield of crop plants (Walter and Kromdijk, 2022). However, better understanding of the mechanisms and consequences of increasing efficiency of photosynthetic reactions in different model species is required to accelerate implementation of these strategies into crops. Overexpression of Rieske previously showed promising results in the model species *A. thaliana* producing higher electron transport and CO_2_ assimilation rates as well as increased biomass accumulation and seed yield (Simkin et al., 2017). Here we tested whether those results could be translated into tobacco, an agronomic crop and a model for other Solanaceae crops, like tomato and potato.

Overexpression of Rieske in tobacco increased the abundance of functional Cyt*b*_6_*f*. Previous works showed that plants overexpressing Rieske had higher leaf content of other subunits of the complex (Simkin et al., 2017) or more Cyt*b*_6_*f* (Ermakova et al., 2019). Here we expand on those results and demonstrate that leaves of Rieske-OE tobacco plants contain significantly more of functionally active Cyt*b*_6_*f* using the CytF activity assay. Whilst the total abundance of Rieske in leaves of transgenic plants was on average double compared to WT, we detected an average of 40% increase of Cyt*b*_6_*f* abundance (Fig. 2). These results suggest that assembly of the complex was likely limited either by an integration of Rieske into the thylakoid membrane (Molik et al., 2001) or by the availability of other subunits. Furthermore, transgenic plants showed only 20% increase of CytF activity compared to control plants (Fig. 3). It is currently thought that Cyt*b*_*6*_*f* assembly starts with the membrane insertion of Cytochrome *b*_6_ and subunit IV and their dimerisation, followed by the insertion of CytF and Rieske (Hasan et al., 2013; Schöttler et al., 2015). Thus, Cyt*b*_*6*_*f* complexes immunodetected with Rieske antibody should already contain CytF (Fig. 2 and Fig. 3). However, it is conceivable that some (virtually) fully assembled complexes are inactive in electron transport, for example, if heme-less CytF is assembled (Mould et al., 2001). Further research is required to identify whether co-overexpression of Rieske with other subunits or regulatory proteins is required to ensure functionality of all assembled complexes.

Rieske-OE plants reached an increased Cyt*b*_6_*f* activity *in vivo*. The first line of evidence came from the LIFT measurements. Plants overexpressing Rieske had more oxidised Q_A_ than WT, meaning that their PSII centers were kept more open after the onset of actinic light (Fig. 5). PQ pool was also more oxidised in Rieske-OE lines, and the rates of electron transport from PSII to PQ pool (Tau1) and from PQ pool to PSI were faster, compared to WT. These results suggested that increasing abundance of Cyt*b*_6_*f*, relative to PSI and PSII (Fig. 4), could accelerate intersystem electron transfer reactions and keep PQ pool transiently more oxidised. The second line of evidence for the increased *in vivo* Cyt*b*_6_*f* activity in Rieske-OE tobacco plants came from the absorbance measurements at 535 nm which reflect both zeaxanthin formation and the modification of the LHCII induced by PsbS (Wilson et al., 2021). Since both components are sensitive to the lumenal pH, the rise of the 535 nm signal in illuminated leaves informs on the kinetics of ΔpH build-up across the thylakoid membrane, largely mediated by Cyt*b*_6_*f*. Three Rieske-OE lines showed a faster rise of 535 nm signal suggesting a faster acidification of the lumen which is consistent with the increased abundance of Cyt*b*_6_*f*. However, *pmf*, representing the total energisation of the thylakoid membranes which is a sum of ΔpH and Δψ (the membrane potential), sampled after 3 min of illumination, did not differ between Rieske-OE and WT plants (Fig. 6). Therefore, the kinetics of the 535 nm signal indicated that electron transport, initially increased in transgenic plants, was subsequently slowed down to a WT-level by the end of 3-min illumination period. Similar trends were observed in the redox states of Q_A_ and PQ pool measured by LIFT: the largest differences between transgenic and WT plants were observed 1-2 minutes after the onset of illumination whilst, towards the end of the 3-minute illumination, all lines were approaching similar values (Fig. 5).

The observed transient increases in Cyt*b*_6_*f* activity were possible because the formation of q_E_ takes time to respond to ΔpH. NPQ is a part of the complex regulatory network that is in place to balance the activity of light and dark reactions of photosynthesis in order to protect the photosynthetic machinery from damaging effects of harvesting excess light energy. When the capacity of light reactions exceeds the utilisation of NADPH and ATP by the Calvin cycle, regulation of ATP synthase conductivity and cyclic electron flow helps to build up the proton gradient, activate NPQ and finally reduce electron transport rate (Kanazawa and Kramer, 2002; Avenson et al., 2005). The conserved relationship between NPQ and *pmf* would constrain the rate of electron transport which is in line with our finding showing that the steady-state electron transport rate was not affected by the increased Cyt*b*_6_*f* activity. Thus, our results indicate that Cyt*b*_6_*f* is not the only factor controlling electron transport rate at high light and high CO_2_ (Fig. 7), the conditions in which electron transport is expected to limit CO_2_ assimilation (Farquhar et al., 1980; von Caemmerer and Farquhar, 1981). In accordance, the increased electron transport and CO_2_ assimilation rates detected in Rieske-OE *A. thaliana* plants could not be solely attributed to the increased Cyt*b*_6_*f* activity, since higher abundances of other photosynthetic complexes were also reported (Simkin et al., 2017).

Photosynthetic control, in response to higher light levels, exerts an additional constraint on the electron transport in Rieske-OE plants. This effect can mask changes in the intrinsic activity of Cyt*b*_6_*f*. As such, changes in Cyt*b*_6_*f* activity is expected to be most readily observed under light conditions the plant is acclimated to i.e. growth light levels. This is exactly what we observe: transient increases in Cyt*b*_6_*f* activity were only clearly resolved at light levels 400 µmol m^-2^ s^-1^ similar to growth conditions (Fig. 5 and Fig. 6). We note that under these conditions, the model of Johnson and Berry can be used to estimate the activity of Cyt*b*_6_*f* and thus the maximal rate of electron transport. The model assumes photosynthetic control is relaxed. That is to say, both NADP^+^ and ADP/P_i_ are available and do not limit Cyt*b*_6_*f* activity. It is important to recognize though that the maximal rate of electron transfer is dependent on the light level used.

At higher light intensities and ambient CO_2_ levels, limited CO_2_ availability slows down the consumption of NADPH and ATP by the Calvin cycle which rapidly downregulates Cyt*b*_6_*f* activity via photosynthetic control before slower regulatory processes like q_E_ and state transitions develop. Although the mechanism of photosynthetic control is not known, it is suggested to be an intrinsic feature of Cyt*b*_*6*_*f* (Malone et al., 2021). It would be interesting to explore whether there is a natural variation for a degree of control this phenomenon exerts over electron transport. Unravelling molecular mechanisms of photosynthetic control would allow optimising it for improving plant productivity.

Due to the above-mentioned limitations, increased Cyt*b*_*6*_*f* abundance and activity in Rieske-OE plants failed to stimulate steady-state electron transport rate in either controlled or field conditions (Fig. 7 and Fig. 8). Consequently, steady-state CO_2_ assimilation rates were not increased in Rieske-OE lines, except for the plants grown in Puerto Rico, which, however, had an elevated CO_2_ availability (Fig. 8). Nevertheless, the detected transient increases in Cyt*b*_*6*_*f* activity suggest that Rieske overexpression could be beneficial if combined with other traits stimulating downstream processes. For example, overexpression of sedoheptulose-1,7-bisphosphatase enzyme of the Calvin cycle is known to stimulate carbon assimilation (Lefebvre et al., 2005; Driever et al., 2017) and could accelerate the use of NADPH and ATP and allow higher electron transport rate in Rieske-OE plants. Simultaneous stimulation of both electron transport and Calvin cycle was previously shown to increase tobacco biomass and yield in the field (López-Calcagno et al., 2020). Testing and combining targets for improving photosynthetic efficiency and deciphering molecular mechanisms of photosynthetic regulation will help to provide necessary improvements to photosynthesis crucial for improving crop productivity.

## Conclusion

Cyt*b*_*6*_*f* is suggested to be one of the main control points limiting the rate of electron transport. Here we generated tobacco plants with increased Rieske content that had more functional Cyt*b*_*6*_*f*, compared to control plants, without marked changes in abundance of other photosynthetic components. Plants with increased Cyt*b*_*6*_*f* abundance displayed a faster build-up of transmembrane proton gradient and higher electron transport rates, however, only transiently. In contrast to Rieske overexpression in Arabidopsis (Simkin et al., 2017), steady-state rates of electron transport and CO_2_ assimilation were not affected in tobacco overexpressing Rieske, pointing out to differences between species. These results suggest that Cyt*b*_*6*_*f* was not the only factor limiting electron transport at high light and high CO_2_ in tobacco.

## Materials and methods

### Generation of construct and transgenic plants

The construct for Rieske overexpression (Fig. S1) was created using the Golden Gate cloning system (Engler et al., 2014). The coding sequence of the *A. thaliana PetC* gene (AT4G03280) was assembled with the *A. thaliana RbcS2B* (*AT5G38420*) promoter and the *A. thaliana HEAT SHOCK PROTEIN 18*.*2* terminator (*AT5G59720*). The construct contained two selection markers: the bialaphos resistance gene (*bar*) under the control of the nopaline synthase promoter (*pNOS*) and the hygromycin phosphotransferase gene (*HPT*) under the control of the cauliflower mosaic virus *35S* promoter. The resulting plasmid was introduced into *N. tabacum* cv. Petit Havana using *Agrobacterium tumefaciens* strain LBA4404 via leaf-disc transformation (Horsch et al., 1985). The shoots were regenerated on Murashige and Skoog medium containing 20 mg L^- 1^ hygromycin and 400 mg L^-1^ cefotaxime. Resistant primary transformants (T_0_ generation) with established roots were transferred to soil and allowed to self-fertilize. T_0_ plants were analysed for the *HPT* copy number (iDNA genetics, Norwich, UK). Homozygous and azygous plants were selected in the T_1_ generation. All experiments were conducted on T_2_ plants. WT plants were used as control for most experiments, except for the field trials where a combination of WT and azygous plants was used.

### RNA isolation and qPCR

Total RNA was extracted from tobacco leaf disks sampled from T_1_ plants and quickly frozen in liquid nitrogen using the NucleoSpin® RNA Plant Kit (Macherey-Nagel, Düren, Germany). cDNA was synthesized using 1 µg of total RNA according to the protocol in the RevertAid Reverse Transcriptase kit (Fermentas, London, UK). Transcript abundance of *AtPetC* and *NtActin* in cDNA samples was estimated using primers AACGCCCAAGGAAGAGTCGT and ACCACCATGGAGCATCACCA for *AtPetC* and CCTGAGGTCCTTTTCCAACCA and GGATTCCGGCAGCTTCCATT for *NtActin* (Schmidt and Delaney, 2010). For qPCR, the SensiFAST SYBR No-ROX Kit was used according to manufacturer’s recommendations (Bioline Reagents Ltd., London, UK).

### Plant growth conditions

For physiological analyses (Fig. 4, Fig. 5, Fig. 6 and Fig. 7), plants were grown from seeds in 2.8-4.5 L pots filled with potting mix and topped with seed raising mix, both supplemented with 7 g L^-1^ of slow-release fertilizer (Osmocote, Scotts, Bella Vista, Australia). Plants were grown in controlled environment rooms at ambient CO_2_, 12 h day /12 h night photoperiod, 24 °C day/20 °C night and 55% relative humidity with light at the intensity of 400 µmol m^−2^ s^−1^ supplied by 1000 W red sunrise 3200 K lamps (Sunmaster Growlamps, Solon, OH). For the thylakoid isolation (Fig. 2 and Fig. 3), plants were grown in cabinets in the same conditions and the illumination was supplied by a mixture of fluorescent tubes (Master TL5 HO 54W/840, Philips Lighting, Roosendaal, The Netherlands) and halogen incandescent globes (42W 2800K warm white clear glass 630 lumens, CLA, Brookvale, Australia). A subset of lines (R17, R20, R26) for the thylakoid isolation was grown in a commercial potting mix supplemented with 2 g L^-1^ of Osmocote; the CytF activity measured from the WT thylakoids grown in different potting mixes did not differ (Fig. S3). All experiments were performed on the first fully expanded leaves (3^rd^ or 4^th^ from the top) of 4-5 weeks old plants.

### Gas exchange

The net rates of CO_2_ assimilation were measured using a portable gas exchange system LI-6800 (LI-COR Biosciences, Lincoln, NE, USA) equipped with the Fluorometer head 6800-01A (LI-COR Biosciences) under red-blue (90%/10%) actinic light. Before the measurements, plants were adapted for 30 minutes to 1500 µmol m^-2^ s^-1^, 400 ppm CO_2_ (reference side), 25°C leaf temperature, 55% humidity, 500 µmol s^−1^ flow rate. The CO_2_ response curves of CO_2_ assimilation rate were measured at stepwise increases of CO_2_ from 0 to 1600 ppm in 3-minute intervals. Multiphase flash of 10,000 µmol m^-2^ s^-1^ (ramp 25%) was applied to leaves in the end of each interval to transiently close all PSII reaction centers and monitor the effective quantum yield of PSII, Y(II) (Genty et al., 1989). The maximum carboxylation rate allowed by Rubisco (*V*_cmax_) and the rate of photosynthetic electron transport (*J*) were obtained by fitting the CO_2_ response curves of CO_2_ assimilation using the routine of Sharkey et al. (2007). *V*_cmax_ was estimated at *C*_i_ < 400 µbar and *J* was estimated at *C*_i_ between 400 and 800 µbar. Mesophyll conductance to CO_2_ of 0.5 mol m^−2^ s^−1^ bar^−1^, previously determined for tobacco (Clarke et al., 2022), was used for the fitting.

### Field experiments

The plants were grown as described in López-Calcagno et al. (2019), and with a methodology broadly analogous to that used commercially for tobacco. Two different field sites were used: Mercedita, Puerto Rico, USA and the University of Illinois Energy Farm, Illinois, USA. The field sites were prepared as described in Kromdijk et al. (2016). The same randomized block design was used in both experiments (Fig. S5). Each block contained all five genotypes in five rows of four plants per genotype. The plants were grown in rows spaced 30 cm apart, with the outer boundary being a WT border. The entire experiment was surrounded by two rows of WT borders. The plants were irrigated when required using rain towers. Seed were germinated in growth cabinets and at 12 d post-germination plants were moved to Trans-plant hydroponic trays (GP009, Speedling Inc., Sun City, FL, USA) and grown in the glasshouse for 20 d before being moved to the field. The plants were allowed to grow in the field until flowering (approximately 30 d). In the field, gas-exchange and fluorescence parameters were recorded using a portable infrared gas analyser LI-6400 (LI-COR Biosciences) with a 6400-40 fluorometer head unit as described in López-Calcagno et al. (2020).

### Electrochromic shift and q_E_

The electrochromic shift signal (ECS) was monitored as the Δ550-515 nm absorbance change and the q_E_ signal was recorded at 535 nm as described in Wilson et al. (2021) using the Dual-PAM-100 (Heinz Walz, Effeltrich, Germany) equipped with the P515/535 emitter-detector module (Heinz Walz) and coupled to the GFS-3000 gas-exchange unit with the Dual-3010 gas-exchange cuvette (Heinz Walz). After 40 min of dark adaptation, a single turnover flash was first applied to record the ECS_ST_ value. After that the leaf was subjected to 3 min light/3 min dark intervals with red actinic light of stepwise increasing irradiance from 60 to 1500 µmol m^−2^ s^−1^. The proton-motive force (*pmf*) was estimated from the amplitude of the rapid ECS decay upon the light-to-dark shift normalised for ECS_ST_. The proton conductivity of the thylakoid membrane (*g*_H+_) was calculated as the inverse time constant of the first-order exponential ECS decay (Kramer and Crofts, 1989).

### Light-induced fluorescence transients

Fast repetition rate (FRR) method was used to analyse electron transport rates from the LIFT measurements (Kolber et al., 1998). We used a commercial terrestrial LIFT-REM fluorometer (https://soliense.com/LIFT_Terrestrial.php) to apply 300 short-excitation flashlets (470 nm, 1.6 µs duration), each separated by a 2.5-µs dark interval, in a saturation sequence to gradually reduce the PSII acceptor side and PQ pool. The excitation power was sufficient to reduce Q_A_ to more than 90% while minimizing electron flow to PQ pool. After this saturation sequence a relaxation sequence of 90 flashlets was applied with exponentially increasing dark intervals to allow oxidation of the PSII acceptors and PQ pool. Four of these fluorescence transients were captured, averaged and fitted with the FRR model at 5 s intervals. Examples of obtained Chl fluorescence transients are shown in Fig. S4. The Q_A_ redox state, the relative oxidation status of the PQ pool and time constants of electron transport, Tau1 (PSII to PQ) and Tau2 (PQ to PSI), were calculated from the transients using the FRR model (https://soliense.com/documents/LIFT-FRR%20assessment%20of%20PQ%20pool%20size.pdf) as described in detail in Osmond et al. (2021).

The leaf was clipped to a custom-made chamber of LI-6400 of approximately 20 cm^2^, which allowed for accurate and reproducible positioning of the leaf with respect to the light sources as well as for control of CO_2_ and humidity (400 ppm CO_2_ in the reference side, 25°C leaf temperature, 55% humidity, 700 µmol s^−1^ flow rate). After 10 minutes of dark adaptation, a leaf was placed in the chamber and measured in darkness, followed by a far-red illumination (735 nm) at 200 μmol m^-2^ s^-1^, monitored with a SKP-216-R 550-750 nm sensor and light meter (Skye Instruments, Llandrindod Wells, UK), from an LED lamp to excite PSI. Red actinic light (627 nm) at 400 μmol m^-2^ s^-1^ was provided by a SL-3500 C LED light source (PSI, Drasov, Czech Republic) and monitored with an LS-C mini-quantum sensor and ULM-500 light meter (Heinz Walz).

### Thylakoid isolation and chlorophyll analysis

Plants were dark-adapted for 12 h prior to thylakoid isolation in order to metabolize excess starch. All steps were performed in minimal light and at 4 °C. 1-2 fully expanded leaves without the midrib were briefly ground in a standard kitchen blender in ice-cold grinding buffer [50 mM 2-(N-morpholino)ethanesulfonic acid (MES)-NaOH, 5 mM MgCl_2_, 10 mM NaCl, 400 mM sorbitol, 0.2% ascorbate, 0.2% bovine serum albumin, 1% polyvinylpolypyrrolidone (PVPP), pH 6.5] supplemented with 0.5 ml L^-1^ protease inhibitor cocktail (Sigma-Aldrich, St Louis, MO, USA). The material was filtered through two layers of Miracloth (Merck Millipore, Burlingtone, MA, USA) and centrifuged at 3000 g and 4 °C for 5 min. The pellet was resuspended in shock buffer (50 mM MES-KOH, pH 6.5). After the centrifugation (3000 g, 4 °C, 5 min) the pellet was resuspended in resuspension buffer (50 mM MES-NaOH, 3.5 mM MgCl_2_, 7 mM NaCl, 500 mM sorbitol, pH 6.5). After the final centrifugation (3000 g, 4 °C, 5 min), the pellet was resuspended in resuspension buffer supplemented with 20% glycerol at approximately 2-4 (mg Chl) mL^-1^. The Chl (*a*+*b*) content was determined in 80% acetone buffered with 2.5 mM of 4-(2-hydroxyethyl)-1-piperazineethanesulfonic acid (pH 7.4) according to Porra et al. (1989).

### Spectroscopic CytF assay

The assay was performed in dim lab light as in Anderson et al. (1997). Briefly, two identical thylakoid samples (1.1 mL each) containing 150 µM Chl (*a*+*b*) were prepared in measuring buffer [50 mM K_2_HPO_4_ pH 6.5, 0.33 M sorbitol, 1 mM MgCl_2_, 1 mM MnCl_2_, 2 mM ethylenediaminetetraacetic acid (EDTA), 1% Triton X-100]. The samples were vortexed and briefly centrifuged to remove starch. The samples were scanned at 580-520 nm in a split-beam spectrophotometer (V-650, Jasco, Tokio, Japan) to determine the baseline (three scans at 0.2 nm interval, 100 nm min^-1^). Subsequently the reference cuvette was supplemented with 0.6 mM Na-ferricyanide and the sample cuvette with 0.6 mM hydroquinone. After 90-s incubation, the samples were scanned again, and the resulting difference spectrum was used for CytF quantification using the reported extinction coefficient 17.7 cm^2^ mmol^-1^ (Bendall et al., 1971). The spectra were corrected for non-linearity of the baseline using the isosbestic points at 542.5nm and 565 nm.

### Blue Native-PAGE, SDS-PAGE and western blotting

Thylakoid complexes were separated using Blue Native-PAGE according to Ermakova et al. (2019). For SDS-PAGE, leaf discs of 0.5 cm^2^ were ground in 0.5 ml of ice-cold sample buffer [100 mM trisaminomethane (Tris)-HCl, pH 7.8, 25 mM NaCl, 20 mM EDTA, 2% w/v sodium dodecyl sulfate (SDS), 10 mM dithiothreitol, 1% w/v PVPP and 2% v/v protease inhibitor cocktail] using glass homogenizers. The leaf extracts were incubated for 10 min at 65 °C and then centrifuged (4 °C, 1 min, 12,000 g). The clear supernatants were supplemented with 4x SDS-loading buffer (200 mM Tris-HCl, pH 6.8, 0.8% w/v SDS, 40% glycerol, 50 mM EDTA, 10% β-mercaptoethanol, 0.08% bromophenol blue), incubated at 65 °C for 10 minutes and loaded on leaf area basis on pre-cast polyacrylamide gels [Nu-PAGE 4–12% Bis-(2-hydroxyethyl)-amino-tris(hydroxymethyl)-methane (Bis-Tris) gel, Invitrogen, Life Technologies Corporation, Carlsbad, CA, USA]. The gels were run in running buffer [50 mM MES, 50 mM Tris, 0.1% w/v SDS, 20mM EDTA, pH 7.3] at a constant voltage of 150 V.

SDS and Blue Native gels were incubated for 10 min in a Bicine-Bis-Tris transfer buffer (25 mM Bicine, 25 mM Bis-Tris, 0.1 mM EDTA, 20% MeOH, pH 7.2) and a Tris-glycine transfer buffer (25 mM Tris, 192 mM glycine, 20% MeOH, pH 8.3), respectively. The proteins were transferred onto a nitrocellulose membrane using a semi-dry system (BioRad, Hercules, CA, USA) according to the manufacturer’s protocol. The membranes were probed with antibodies against various photosynthetic proteins in a dilution recommended by the producer: Rieske (AS08 330, Agrisera, Vännäs, Sweden), CytF (Price et al., 1995), PC (Agrisera, AS06 141), PsaB (Agrisera, AS10 695), D1 (Agrisera, AS10 704), AtpB (Agrisera, AS05 085), Lhcb2 (Agrisera, AS01 003), PsbS (Agrisera, AS09 533), RbcL (Martin-Avila et al., 2020). Secondary goat anti-rabbit horse radish peroxidase-conjugated antibody (Biorad) was detected using Western lightning Ultra chemiluminescence kit (Perking Elmer, Waltham, MA, USA). Quantification of the signals was performed using ImageJ software.

### Statistical analysis

Type II ANOVA or two-tailed, heteroscedastic Student’s *t*-test were performed to test the relationship between mean values for transgenic and control plants. Details of tests and replicates are provided in figure legends.

## Acknowledgements

This study was funded by the Bill and Melinda Gates Foundation RIPE (Realizing Increased Photosynthetic Efficiency) programme. The Australian Plant Phenomics Facility supported under the National Collaborative Research Infrastructure Strategy of the Australian Government is gratefully acknowledged for the use of growth facilities and equipment. We thank Dean Price and Spencer Whitney for the gift of antibodies. We thank Z. Kolber for frequent and tireless help with LIFT.

## Supplementary materials

**Fig. S1.**
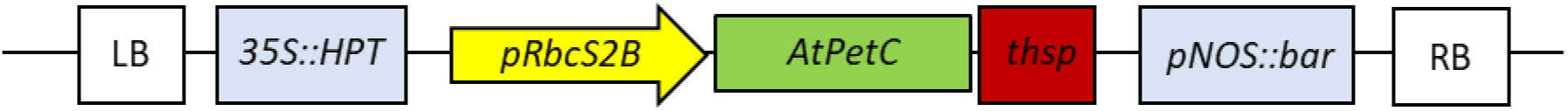
Schematic representation of the gene construct used for Rieske overexpression. LB, T-DNA left border; *35S*, cauliflower mosaic virus *35S* promoter; *HPT*, hygromycin phosphotransferase gene; *pRbcS2B, A. thaliana* Rubisco small subunit 2B (*AT5G38420*) promoter; *AtPetC, A. thaliana PetC* (AT4G03280) coding sequence; *thsp, A. thaliana HEAT SHOCK PROTEIN 18*.*2* (*AT5G59720*) terminator; *pNOS*, nopaline synthase promoter; *bar*, bialaphos resistance gene; RB, T-DNA right border.

**Fig. S2.**
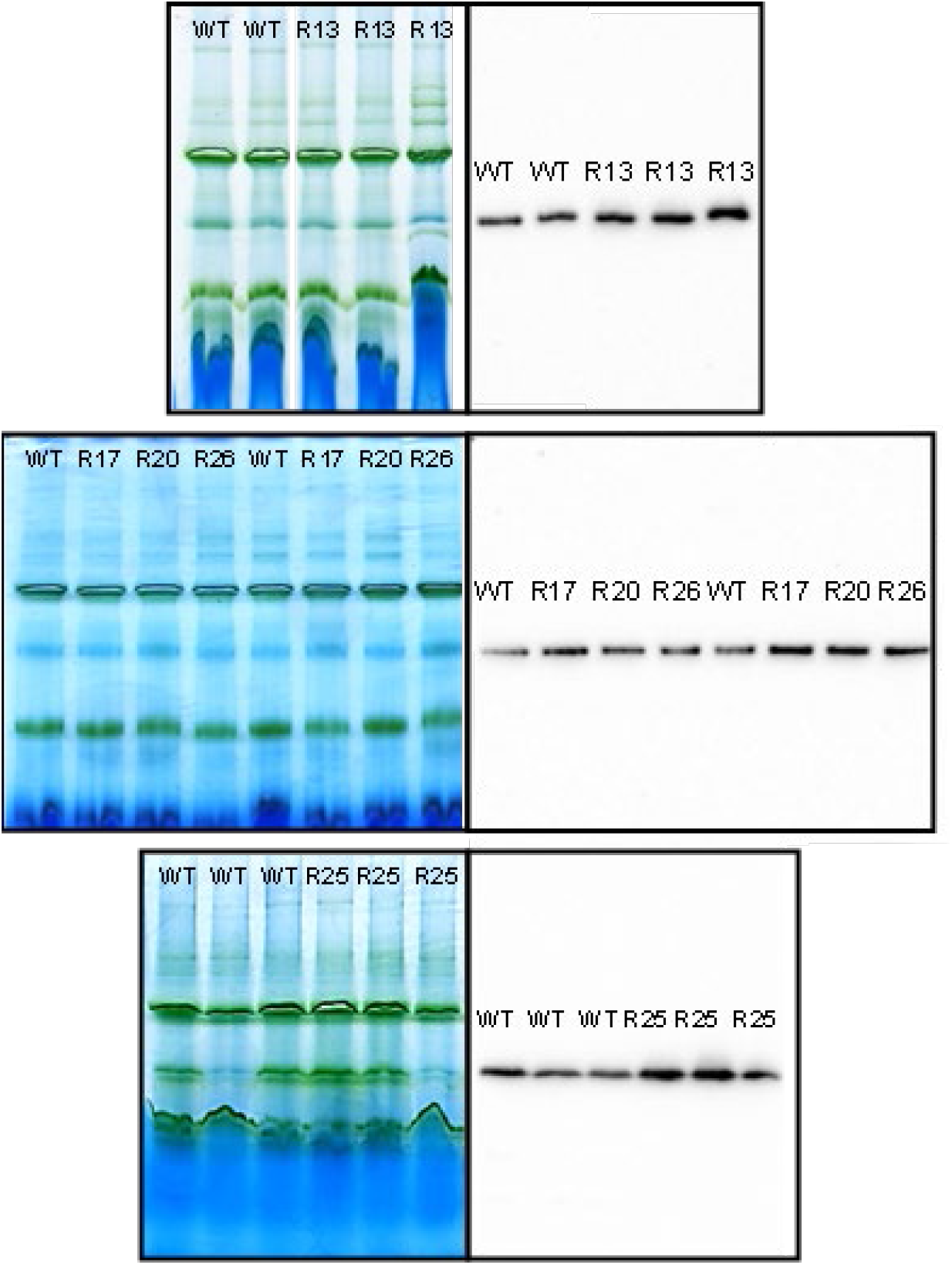
BN-PAGE and blots performed in this study. Relative quantification of western blot signals is shown in Fig 2.

**Fig S3.**
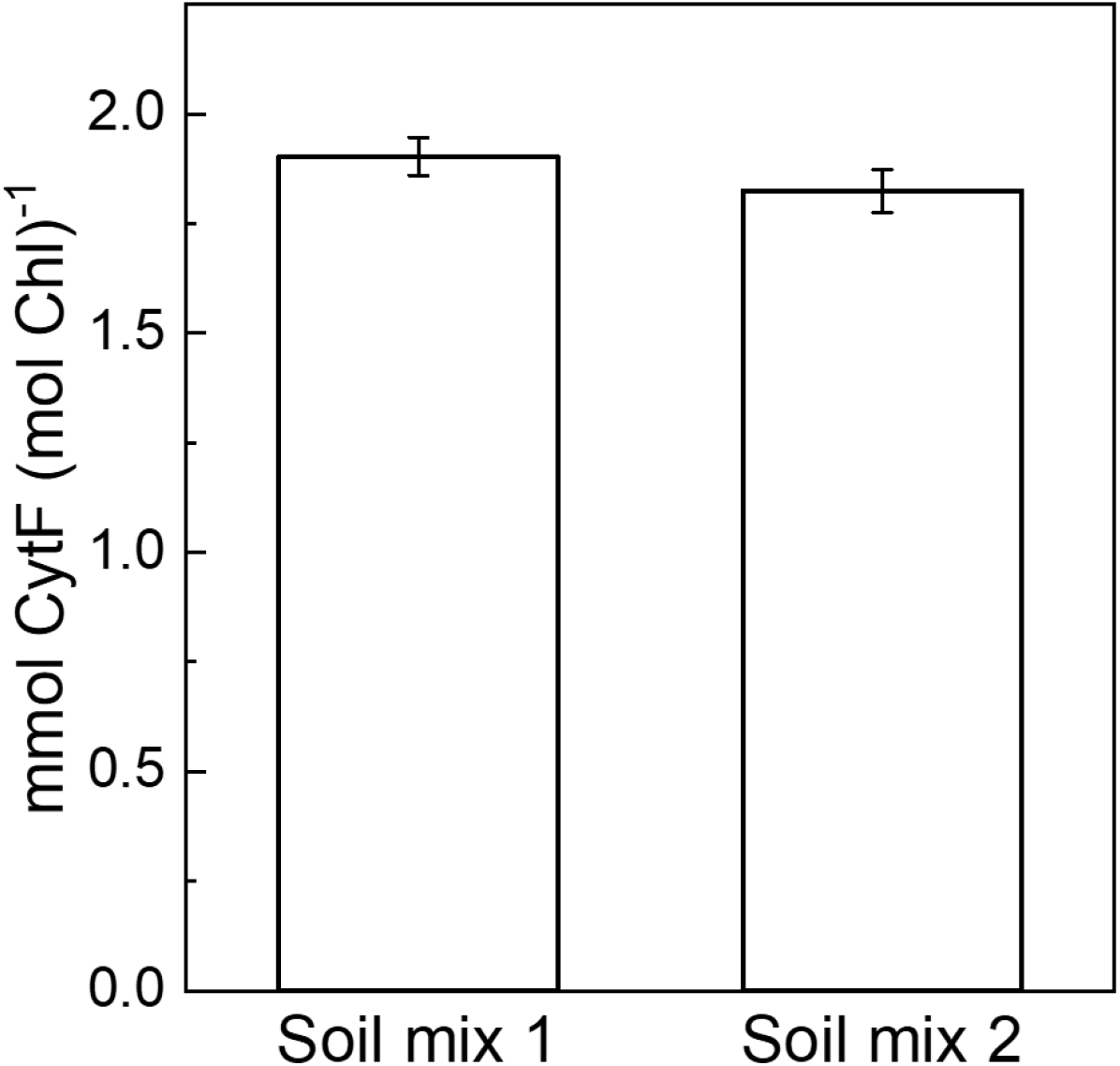
Comparison of the thylakoid CytF content in plants grown in a commercial soil mix supplemented with 2 g L^-1^ osmocote (mix 1) and a homemade soil mix supplemented with 7 g L^-1^ osmocote (mix 2). Mean ± SE, *n* = 3 biological replicates for mix 1, *n* = 5 for mix 2. Not significant (*t*-test, *P* = 0.63).

**Fig. S4.**
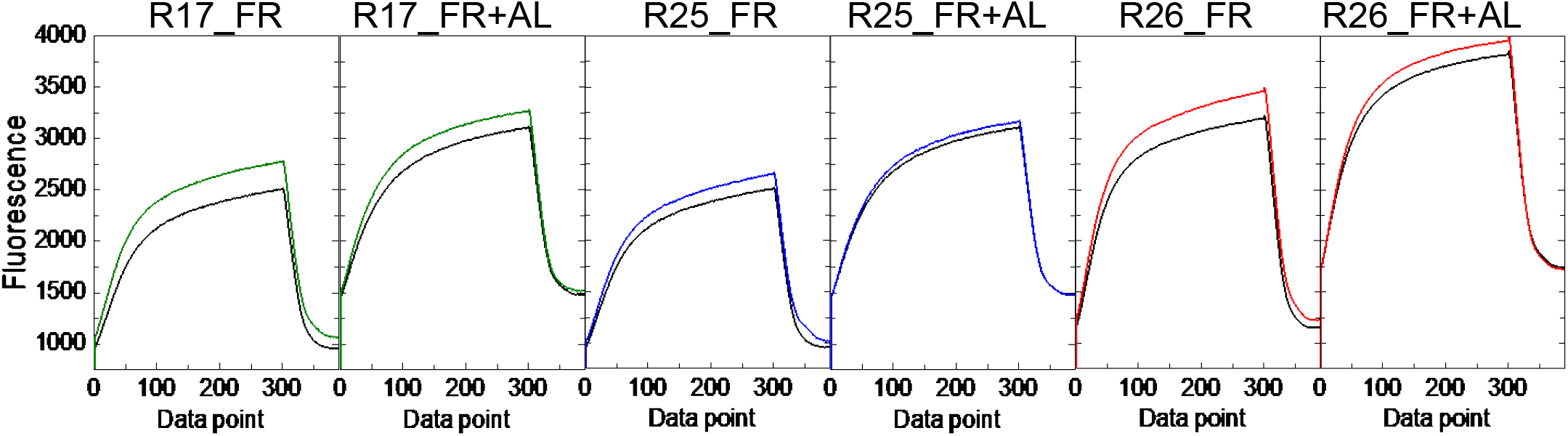
Fluorescent transients obtained by LIFT analysis from WT (black traces) and Rieske-OE plants of lines R17, R25 and R26. Two LIFT-FRR duty cycles are presented for each line. FR, the last point at the end of far-red illumination, prior to switching on the actinic light; FR+AL, the point measured after 3 minutes of illumination with far-red and actinic light.

**Fig. S5.**
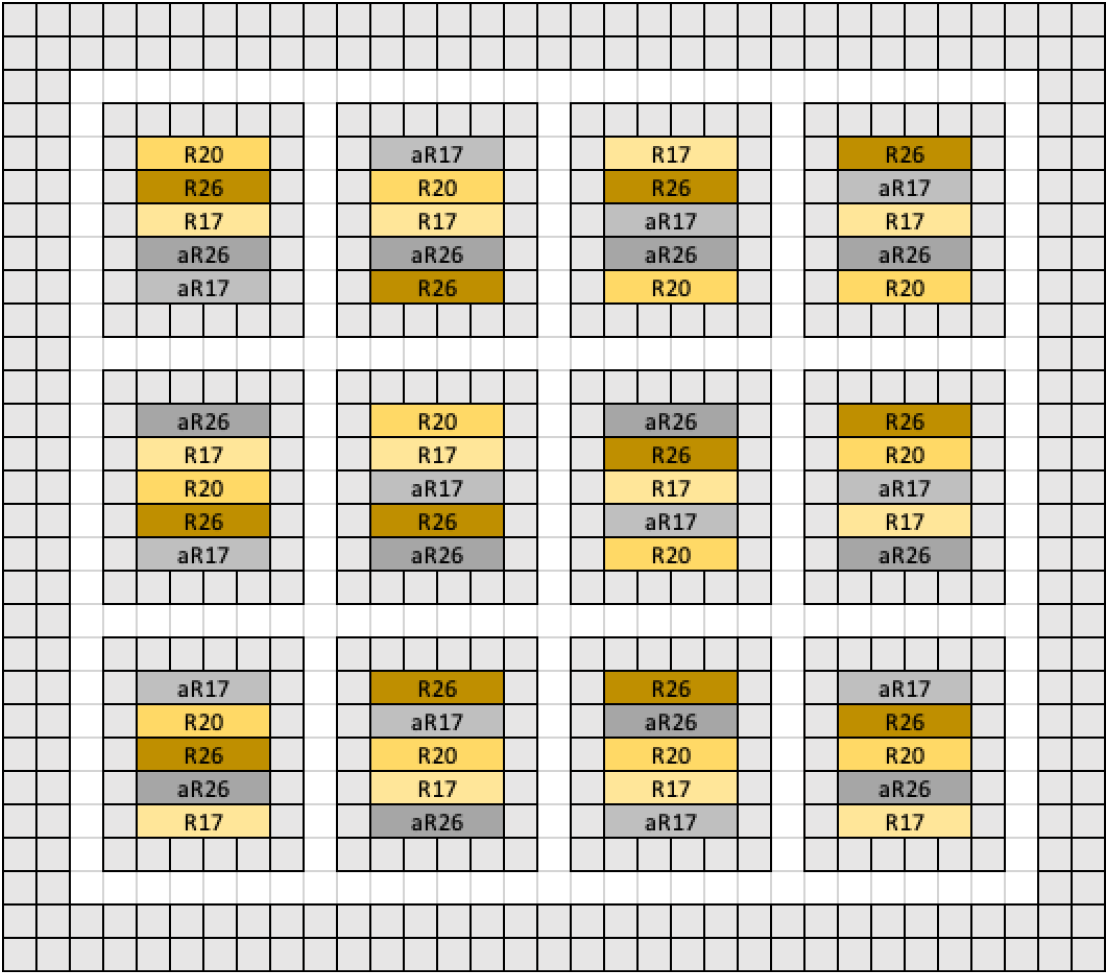
Schematic representation of experimental design for the field experiments. A randomized complete block design was used for both experiments where each block had 4 plants for every line. Rows were randomised using RAND function (Microsoft Excel 2010). Both experiments were surrounded by a WT border and each separate experiment was bordered by two lines of WT.

**Fig. S6.**
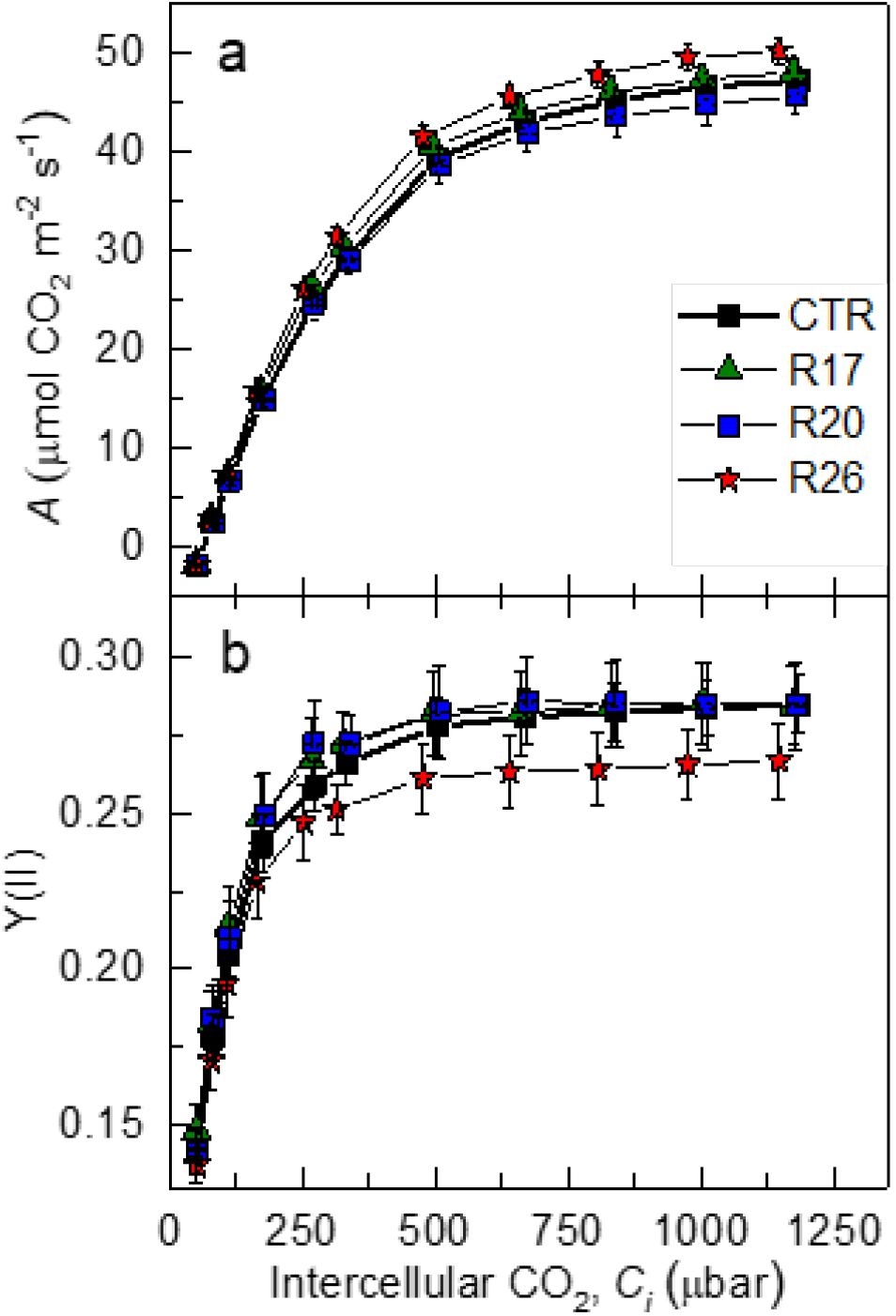
CO_2_ response of (**a**) CO_2_ assimilation, *A*, and (**b**) the effective quantum yield of PSII, Y(II), of Illinois field-grown control and Rieske-OE plants measured at 2000 µmol m^-2^ s^-1^. The control group (CTR) represents both WT and azygous plants. Mean ± SE, *n* = 18 for CTR plants, *n* = 12 for transgenic lines. No significant differences were found between the genotypes (a linear mixed-effects model and type III ANOVA, *P* < 0.05).

